# Phagocytosed polyhedrin-cytokine co-crystal nanoparticles provide sustained secretion of bioactive cytokines from macrophages

**DOI:** 10.1101/2021.02.17.431660

**Authors:** Astrid Wendler, Nicholas James, Michael H Jones, Christian Pernstich

## Abstract

Many cells possess the ability to engulf and incorporate particles by phagocytosis. This active process is characteristic of microorganisms as well as higher order species. In mammals, monocytes, macrophages and microglia are among so-called professional phagocytes. In addition, cells such as fibroblast and chondrocytes are classified as non-professional phagocytes. Professional phagocytes play important roles in both the innate and adaptive immune response, wound healing and tissue homeostasis. Consequently, these cells are increasingly studied as targets and vectors of therapeutic intervention to treat a range of diseases. Professional phagocytes are notoriously difficult to transfect limiting their study and manipulation. Consequently, efforts have shifted towards the development of nanoparticles to deliver a cargo to phagocytic cells via phagocytosis. However, this approach carries significant technical challenges, particularly for protein cargos. We have focused on the development of nanoscale co-crystalline protein depots, known as PODS®, that contain protein cargos, including cytokines. Here, we show that PODS are readily phagocytosed by non-professional as well as professional phagocytic cells and have attributes, such as highly sustained release of cargo, that suggest potential utility for the study and exploitation of phagocytic cells for drug delivery. Monocytes and macrophages that ingest PODS retain normal characteristics including a robust chemotactic response. Moreover, the PODS-cytokine cargo is secreted by the loaded cell at a level sufficient to modulate the behavior of surrounding non-phagocytic cells. The results presented here demonstrate the potential of PODS nanoparticles as a novel molecular tool for the study and manipulation of phagocytic cells and for the development of Trojan horse immunotherapy strategies to treat cancer and other diseases.

## Introduction

Phagocytic cells engulf microscopic particles through an active process (1) and are classified as professional or non-professional with the latter taking up a more limited range of materials. Professional phagocytes include monocytes, macrophages, microglia, dendritic cells and neutrophils. Non-professional phagocytes include chondrocytes, fibroblasts, erythrocytes and epithelial cells.

Macrophages contribute to the innate and adaptive immune response. They also help to heal wounds and regulate tissue homeostasis(2). Macrophages display phenotypic plasticity and can be polarized by microenvironmental cues to a spectrum of phenotypes very broadly characterized by the classically activated pro-inflammatory M1 phenotype and the alternatively activated anti-inflammatory M2 phenotype. Macrophages are able to switch between these different phenotypes depending on changing requirements (3).

The role of macrophages in cancer underscores their importance. In pre-cancerous lesions, macrophages target and remove defective cells, but as cancers develop, immuno-editing generates cancer cells that avoid this surveillance. Most tumor associated macrophages (TAMs) are derived from monocytes which are constantly recruited into malignant tumors by chemotactic signals generated by the tumor, such as the CCL2 chemokine axis (4,5). Once inside the cancer, the monocytes differentiate to macrophages. In later stages of disease, these macrophages are subverted by the cancer cells towards a M2-like phenotype, which actively supports the maintenance of cancer and suppresses the anti-cancer activity of other immune cells (6). As a result of this central role, macrophages, and the proteins they secrete, are an important immunotherapeutic target for cancer drugs. Their tumor infiltrating behavior also suggests their potential as vectors for targeted drug delivery.

The basic study of macrophages and the development of macrophage-based cellular therapies has been hampered by poor nucleic acid transfection efficiency (7). This challenge has led many to explore the utility of nanoparticles carrying a cargo of functional molecules. To be useful, nanoparticles must be efficiently phagocytosed and release their cargo intact from the phagolysosome. The cargo must then be transported to targets within the cell or secreted by the cell to engage with extracellular targets.

A wide range of nanoparticles have been evaluated to date. These include drug-coated nanoparticles (8), radiosensitizers (9), drug-filled synthetic lipid particles (10), enzymes (11), and glucocorticoid pro-drugs (12). These nanoparticles have achieved limited success and a nanoparticle that can usefully deliver proteins, and particularly cytokines, to macrophages has yet to be developed.

Cytokines are key signaling molecules of the immune response. Recombinant cytokines such as IL-2, administered therapeutically, are able to activate immune cells and have been approved to treat certain metastatic cancers. Although high dose IL-2 therapy (Proleukin®/Aldesleukin) is highly effective in some patients, its acutely toxic side effects limit use as a frontline therapy(13). Effective cytokine nanoparticles containing IL-2 and other cargos may allow the development of a macrophage-mediated molecular Trojan horse strategy for the targeted delivery of cytokines to disseminated cancer.

PODS are nanoscale (200 nm-5 µm) protein co-crystals built from the polyhedrin protein. Their distinctive cubic structure is determined by intracellular assembly of a specific polyhedrin protein derived from the *Bombyx mori* cypovirus (14). A cargo protein can be incorporated into PODS during crystal assembly via an immobilization tag, typically using a fragment of the polyhedrin H1 finger. PODS happen to be the ideal size, shape and rigidity for phagocytic uptake (15). PODS are also non-brittle and temperature-stable. In the presence of proteases, PODS slowly degrade and typically release their cargo over a period of one-two months. Fortuitously, this degradation results in the release of cargo protein that retains bioactivity.

Here, we explore the potential of PODS for delivering functional cytokines to monocytes and macrophages. We show that PODS particles are very efficiently phagocytosed and that cargo cytokines either evade or withstand the harsh conditions of the phagolysosome allowing secretion by macrophages. Moreover, secretion levels achieved are sufficient to produce distinct phenotypic changes in heterogenous co-cultured cells. Our findings demonstrate the utility of PODS for macrophage-based research and suggest their potential utility in Trojan horse drug delivery strategies to treat cancer and other diseases.

## Results

### Efficient phagocytosis of PODS crystals into macrophages

PODS protein crystals form near-perfect cubes. Their angular shape and overall appearance make them distinct from cells and thus easily identifiable even at low magnification (Supplementary Figure 1). We first tested our expectation that PODS crystals can be phagocytosed by primary murine monocytes and macrophages. Murine bone marrow derived monocytes (BMDM) were isolated according to Wagner et al (16) from the tibias of C57BL/6 mice. Cells were cultured for 5 days before they were incubated with fresh complete medium containing PODS M-CSF or PODS GM-CSF at a ratio of 5 crystals per cell. PODS crystals were phagocytosed into the BMDMs with high efficiency (Figure 1a), seemingly without negatively influencing their behavior (supplementary material, video 1). Moreover, when PODS GM-CSF-loaded BMDMs were incubated with a further 5 PODS per cell, they ingested all the additional PODS crystals (Supplementary Figure 2) whilst maintaining regular motility and morphological changes, such as protruding leading edges and uropods (Supplementary material, video 2).

**Figure 1:**
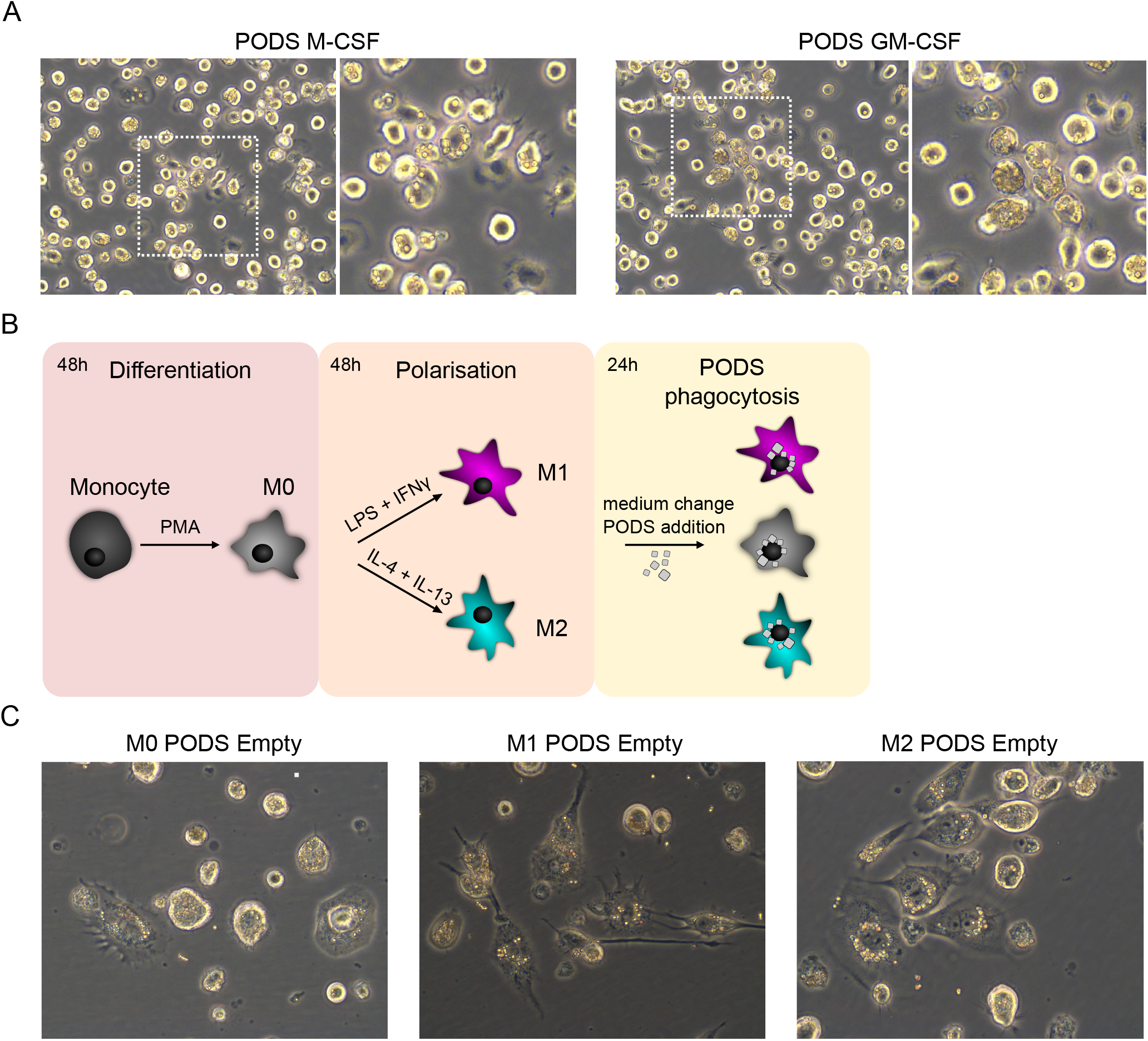
Phagocytosis of PODS protein crystals into professional phagocytes. A) Brightfield images of PODS M-CSF and GM-CSF loaded bone marrow-derived mouse macrophages at a magnification of 40x. The dashed square marks the area of the zoom shown in the right image of each panel. B) Schematic overview of the THP-1 differentiation (M0), polarization (M1, M2) and PODS phagocytosis workflow. C) THP-1 cells were differentiated first into M0 macrophages and then further polarized into either M1 or M2 macrophages according to the workflow depicted in B. Cells of the various differentiated types were cultured with PODS Empty at a concentration of 10 PODS per cell for 24 h and subsequently imaged at a magnification of 40x.

Following successful phagocytosis into primary phagocytes, we used THP-1 cells as *in vitro* model for macrophages, based on Genin et al (17). These cells can be readily polarized to different macrophage types. Figure 1b shows a schematic of the workflow. THP-1 monocytes were differentiated into M0 macrophages by incubating cells in complete medium supplemented with phorbol-12-myristate-13-acetate (PMA). M0 cells were then polarized to M1 or M2 type macrophages by incubating cells in complete medium supplemented with either IFN-γ and LPS or IL-4 and IL-13, respectively. Once polarized, the medium was replaced with fresh complete medium containing PODS Empty crystals (polyhedrin-only, containing no cargo protein) in a ratio of 10 crystals per cell and uptake of PODS was monitored repeatedly over 24 h using a live cell imaging system. Brightfield images taken after 24 h show that almost all crystals have been taken up by the cells, independent of their polarization status (Figure 1c and Supplementary Figure 3). Moreover, non-professional phagocytes such as chondrocytes, NIH-3T3 fibroblasts and C2C12 myoblasts, were equally capable of ingesting PODS efficiently (Supplementary Figure 4).

### Viability and polarization status of PODS-loaded macrophages

Given the cytotoxic effect generated by different types of nanoparticles on macrophages in previous studies (18–20), the viability of M0, M1 and M2 cells was examined at 48 h (Figure 2a) and 96 h (Supplementary Figure 5) after uptake of different numbers of PODS crystals. Results from WST-8 assays demonstrated that cell viability was not reduced compared with untreated cells (M0, M1 and M2 cells which were not loaded with PODS), suggesting that the PODS particles were non-toxic at a median dosage range of up to 15 PODS/cell. Neither PODS Empty nor PODS FGF-10 (a growth factor which alone is non-toxic to macrophages) (21) had any negative effect on macrophage viability. However, there was a limit to how many PODS a macrophage could ingest before going into apoptosis. Ingestion of around 50 PODS per cell clearly lead to some cells “bursting” into apoptotic bodies (Supplementary figure 6).

**Figure 2:**
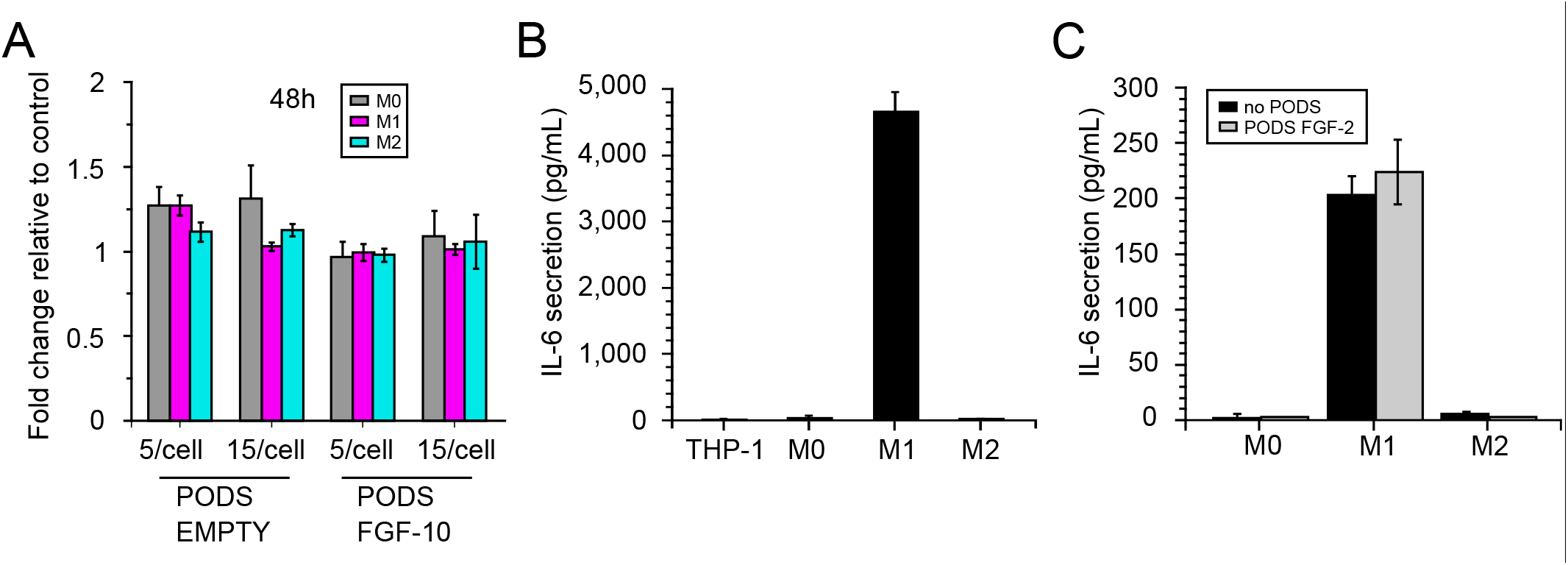
Characterization of professional phagocytes post-phagocytosis of PODS proteins. A) M0, M1 and M2 macrophages were incubated with PODS Empty or PODS FGF-10 (1:5 and 1:15) for 24 h in a 96-well TC plate. Subsequently, medium was changed and viability of cells was measured 48h after PODS uptake using a colorimetric assay (Orangu™, Cell Guidance Systems). The fold change in viability was calculated relative to unloaded macrophages. B) Medium of THP-1, M0, M1 and M2 was collected directly after polarization and tested for the presence of IL-6 by ELISA. C) The same cells were then washed and incubated with PODS FGF-2 for 24 h in fresh full growth medium. The medium was then collected and analyzed for the presence of IL-6 by ELISA.

The activation and polarization of monocytes towards an M1-like phenotype upon contact with a foreign body is a key element of the innate immune system. To determine if uptake of PODS crystals by monocytes and macrophages influences their polarization status, we measured the secretion of IL-6, a marker for M1 polarization, before and after PODS uptake. First, we confirmed that IL-6 can indeed be used as a proxy for polarization status, showing that M1 cells raised concentration levels of IL-6 in the cell culture media to more than 4 ng/ml directly after polarization (Figure 2b), whereas neither non-activated THP-1 cells nor M0 and M2 cells raised IL-6 concentration above background level.

The polarization medium was changed to standard media with no growth factors. The cells were then divided into two groups. In the first group, PODS FGF-2 were added and phagocytic uptake was allowed to proceed for 24 h to load the cells. Following this incubation period, both loaded and unloaded M1 cells secreted IL-6 further, up to concentrations of 200 pg/ml (Figure 2c). The continuous secretion of IL-6 from PODS loaded M1 cells at the same level as unloaded M1 cells suggests that the uptake of PODS crystals did not alter their polarization status. Equally, the uptake of PODS crystals did not induce IL-6 secretion of M0 and M2 cells, suggesting that phagocytosis of PODS crystals by itself does not change the polarization status of any macrophage type.

### PODS-loaded macrophages retain functionality

Having established that PODS particles are efficiently taken up and do not affect macrophage viability, we assessed the cells for their characteristic ability to migrate and follow chemotactic signals.

To quantify potential changes in mobility due to uptake of PODS, M0 cells were incubated with PODS IL-2 for 24 h and then observed with a live cell imaging system under normal growth conditions for a further 24 h. Movement of unloaded- and PODS IL-2-loaded cells was analyzed by tracking each of 25 cells across 720 frames taken at 2-minute intervals (Figure 3a). When comparing tracks for both the distance migrated and randomness, there was no detectable difference in the mobility of cells with and without PODS.

To test the ability of PODS IL-2-loaded M0 cells to follow a chemotactic gradient, the above experiment was repeated with a slight variation: after PODS uptake, cells were taken into serum-free medium and transferred into the observation area of chemotactic slides (<u>μ</u>-Slide, Ibidi) in between two reservoirs. Reservoir 1 was filled with medium containing 10% serum, acting as a chemoattractant, and reservoir 2 was filled with serum-free medium. The movement of unloaded- and PODS IL-2-loaded cells was investigated by tracking 26 cells for 12 h (1 frame per minute) each as described above (Figure 3b). The analysis of this tracking data revealed that uptake of PODS crystals does not impair migration of macrophages towards a chemoattractant as both unloaded and PODS-loaded M0 cells followed the chemical cue to the same extent.

**Figure 3:**
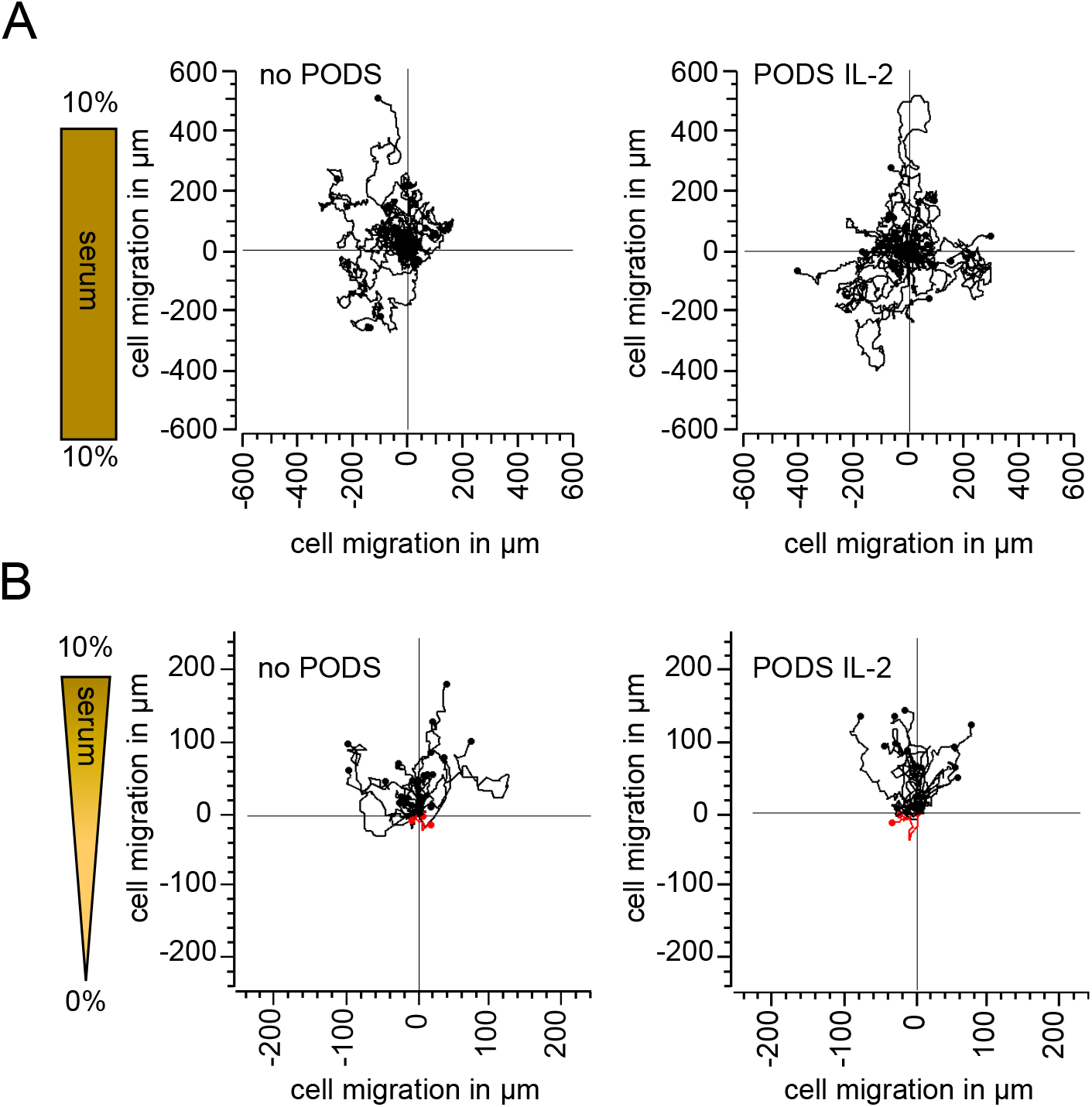
Mobility & chemotaxis of M0 cells loaded up with PODS. A) Mobility of PODS IL-2 loaded macrophages. The mobility of empty and PODS IL-2 loaded macrophages (10 PODS /cell) in full growth medium was monitored using a live cell imaging system. Images were taken every 2 min for 24h and analyzed using the manual tracker of Image J, and the chemotaxis and migration tool (Ibidi). 25 cells per condition were followed and their migration tracks visualized as a rose plot, in which each line represents the track of an individual cell over 24 hours. Track positions were normalized so that each track starts at the (0,0) coordinate. B) Empty and PODS IL-2 loaded macrophages were transferred into Chemotaxis µ-Slides (Ibidi) and tested for their ability to follow a chemoattractant gradient generated from 0-10% BCS. Images were taken every 1 min for 12 h and analyzed using the manual tracker utility of Image J, and the chemotaxis and migration tool (Ibidi). 26 cells per condition were followed and their migration tracks visualized as a rose plot, in which each line represents the track of an individual cell over 12 hours. Track positions were normalized so that each track starts at the (0,0) coordinate. Tracks of cells migrated towards the 10% BCS source are shown in black (up), whereas track of cells migrated away from the BCS source are shown in red (down).

Another characteristic of macrophages is their ability to traverse narrow capillaries and to extravasate into surrounding tissues. To assess if PODS-loaded macrophages retain the ability to traverse narrow spaces, PODS eGFP (enhanced green fluorescent protein)-loaded M0 cells were placed into serum-free medium and transferred to 24-well tissue culture inserts containing extended 8 µm diameter pores, akin to the narrowest blood capillaries, in their bases. Maintenance medium containing 10% serum was used as a chemoattractant in the lower well (Figure 4a). After 24 h of incubation, the number of cells that migrated towards serum was compared to the number of cells in a second chamber which lacked serum as chemoattractant. Significantly larger cell numbers migrated into the wells with chemoattractant demonstrating that loaded M0 cells were capable of passing through the narrow pores and actively migrated through the 8 µm pores in the presence of a chemoattractant.

**Figure 4:**
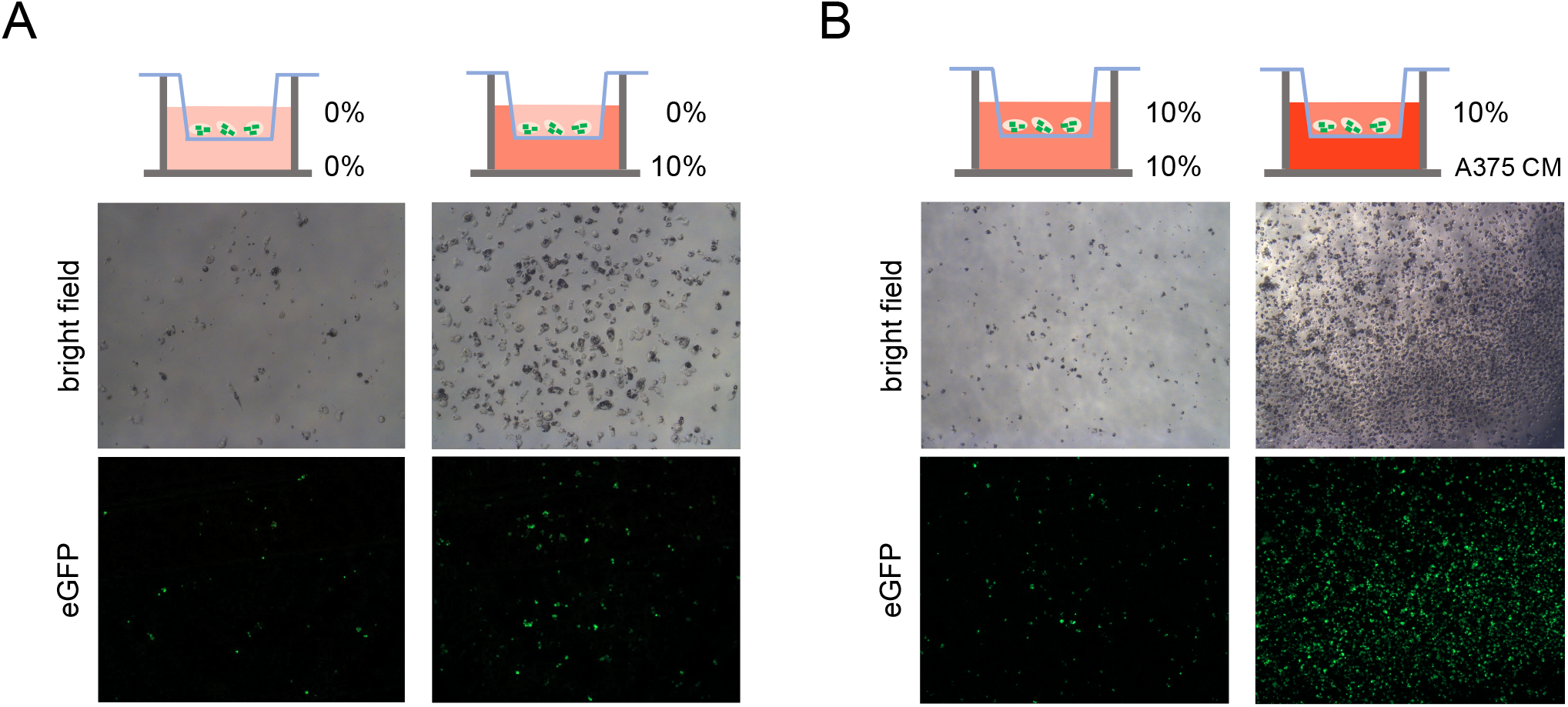
Directed migration of PODS eGFP loaded macrophages through 8 µm pores. A) PODS eGFP loaded macrophages (5 PODS /cell) in serum-free medium were transferred into 24-well inserts with 8 µm pores. The bottom well was filled with either RPMI-1640 only (left) or RPMI-1640 supplemented with 10% FBS (right). The top images show to the bottom well migrated cells in the brightfield channel and the lower images show the corresponding fluorescent channel after 24h of incubation. B) PODS eGFP loaded macrophages follow chemical signals from melanoma cell line through 8 µm pores. PODS eGFP loaded macrophages (5 PODS /cell) in full growth medium were transferred into 24-well inserts with 8 µm pores. The well was filled with either full growth medium (left) or medium that was conditioned with A375 cells for 3 days (right). The top images show to the bottom well migrated cells in the brightfield channel and the lower images show the corresponding fluorescent channel after 24h of incubation.

We are interested in the potential of PODS-loaded macrophages to provide a delivery mechanism to modulate the tumor microenvironment. Macrophages are actively recruited by chemoattractants, such as CCL2, secreted by solid tumors (6). To explore whether PODS-loaded macrophages responded to chemotactic cues secreted by cancer cells, we use conditioned medium from a cancer cell line to attract PODS eGFP loaded macrophages. The same cell chamber described above was used with the PODS loaded cells in the insert and the conditioned media below. To condition the media, A375 cells (a human malignant melanoma cell line) were grown for 3 days in 10% serum containing growth medium. The conditioned medium was then used as chemoattractant in the bottom well (Figure 4b) with the PODS-loaded macrophages placed in the top chamber in media containing 10% serum. As a control, a second cell chamber was set up without conditioned media, containing the same 10% serum in the lower well and upper insert. The cells were incubated for 24 h and then phase contrast as well as fluorescent microscopy were used to observe the number of cells that had moved into the well from the insert. This experiment clearly showed that PODS eGFP-loaded M0 cells were attracted specifically by the cancer cell line-conditioned medium, migrating through the 8 µm pores towards the chemotactic source.

### Release of cargo from PODS-loaded macrophages

Macrophages orchestrate immune responses by secreting a range of cytokines and other signaling molecules which affect other immune cells. We wished to determine if it is possible to modulate this secretion profile using PODS. To test if ingested PODS cargo protein is released from the macrophages into the medium, M0 cells were incubated with PODS IL-6 for 24 h, washed twice and then incubated for a further 4 days in normal growth medium containing no additional PODS. Medium conditioned by the loaded macrophages was collected and analyzed for the presence of IL-6 by ELISA (Figure 5a). Against expectations, cargo protein IL-6 was readily detected in the cell culture medium. Moreover, the levels of IL-6 were dose-dependent: the more PODS IL-6 were loaded into M0 cells, the more IL-6 could be detected in the medium after 4 days. As a control, we also measured the amount of IL-6 released from similar numbers of non-phagocytosed PODS IL-6 (5b). The amount of IL-6 released from naked PODS was 3 -10 times higher than from PODS taken up by macrophages suggesting either reduced rates of cytokine release from macrophages or increased rates of degradation. As a further control, M0 cells were incubated with PODS FGF-2 in the same way and levels of IL-6 in the medium were analyzed IL-6 levels remained below background (<10 pg/ml) (Figure 5a).

**Figure 5:**
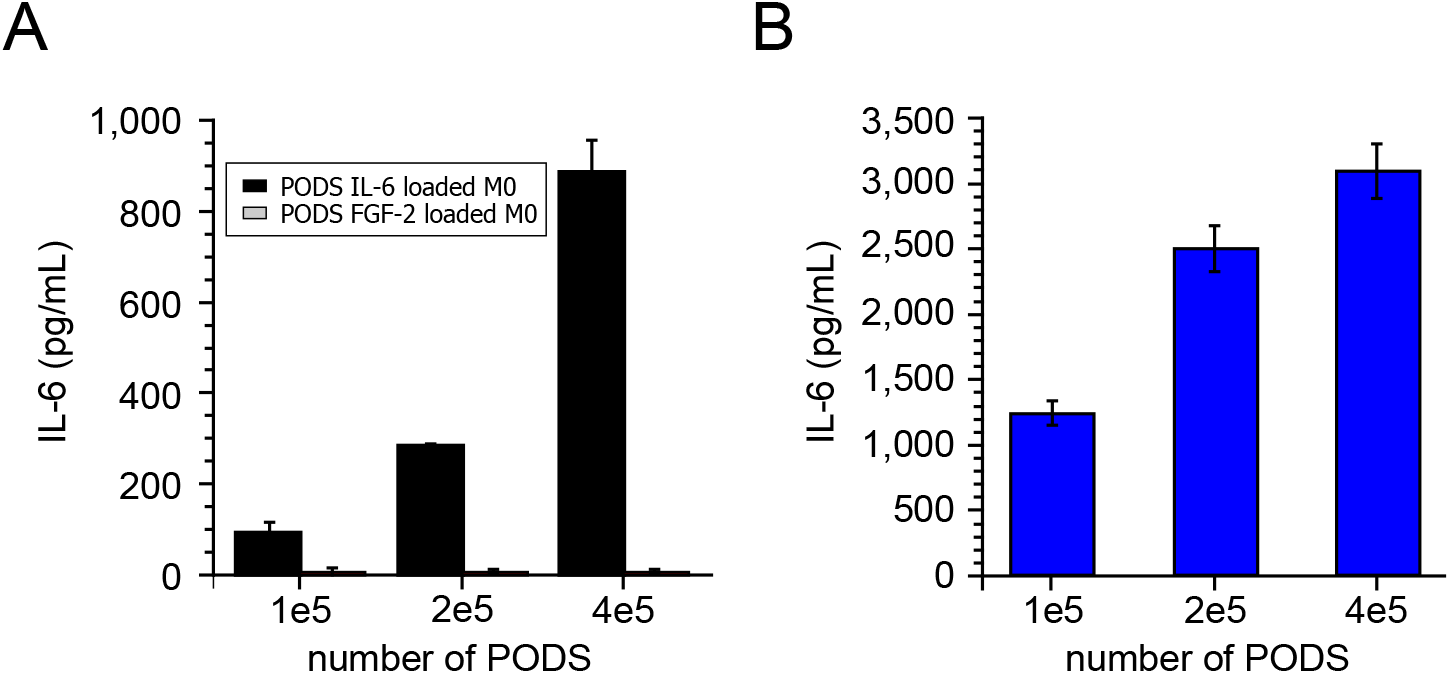
Release of IL-6 from PODS IL-6 loaded macrophages. A) M0 were loaded with 5, 10 or 20 of either PODS IL-6 or PODS FGF-2 per cell (equivalent to 1x 10^5^, 2x 10^5^ and 4x 10^5^ PODS per 96-well). Cells were washed and incubated in full growth medium for 4 d. Medium was collected and tested for the presence of IL-6 in the medium. B) The same amounts of PODS IL-6 as before were spun down to the bottom of a 96-well plate and incubated for 4 d in full growth medium. The medium was collected and levels of IL-6 measured by ELISA.

### Cargo released from PODS-loaded macrophages effects phenotypic change in heterogenous co-cultured cells

After demonstrating secretion of cytokine cargo by macrophages, we wished to determine if release rates are sufficient to drive changes in phenotype in co-cultured heterogenous cells. Here, a proliferation assay using NIH-3T3 cells that are responsive to the FGF-2 protein was performed. NIH-3T3 cells were seeded in normal growth medium into wells of a 24-well plate. After 24 h, the media was replaced with serum-free medium. Separately, M0, M1 and M2 polarized macrophages were incubated with PODS FGF-2 (10 PODS/cell) for 24 h to facilitate phagocytosis. PODS FGF-2-loaded macrophages were then detached, changed into serum-free medium and seeded into 24-well TC inserts which were subsequently placed into the wells that contained NIH-3T3 cells. Co-cultures were incubated for a further 4 days and the number of NIH-3T3 cells assessed by performing a WST-8 assay (Figure 6). Co-incubation of all types of PODS FGF-2-loaded macrophages supported the growth of FGF-2 responsive NIH-3T3 cells compared to unloaded macrophages.

**Figure 6:**
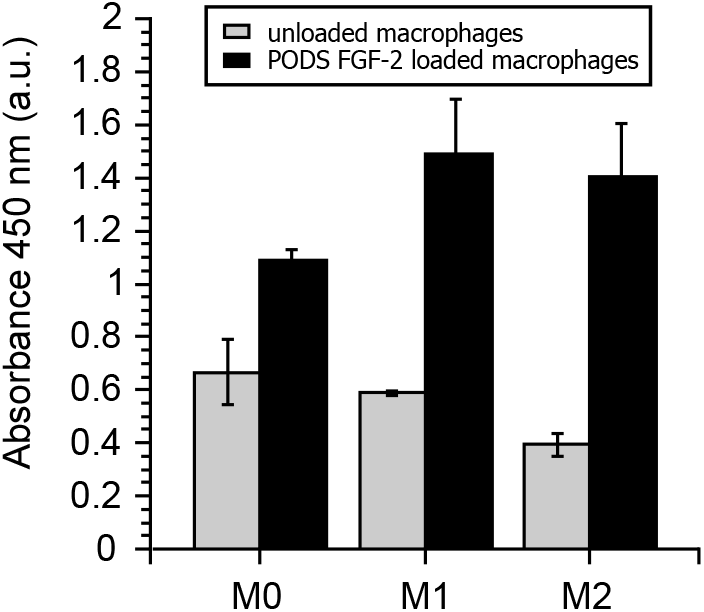
Released cargo from phagocytosed PODS is bioactive. Bioactivity of FGF-2 released from PODS FGF-2 loaded macrophages. Unloaded and FGF-2 loaded macrophages (M0, M1 and M2 with 10 PODS/cell) in TC inserts were incubated with FGF-2 reactive NIH-3T3 cells under serum-free conditions for 4 d. Proliferation of NIH-3T3 cells was measured using a colorimetric assay (Orangu™, Cell Guidance Systems).

### Material & Methods

#### PODS crystal synthesis

All PODS proteins were synthesized as previous described (22,23). All constructs were fused to the H1 immobilization tag (24). Briefly, baculovirus (BV) DNA and transfer DNA was co-transfected into standard *Spodoptera frugiperda* 9 (Sf9) cells using TransIT-Insect (Mirus Bio). Resulting infective BV was harvested and a plaque purification then performed to isolate a single recombinant BV. Isolated plaques were first screened and positive BV then harvested, expanded and finally used to infect large scale Sf9 cells cultures to produce PODS crystals. Subsequently, crystals were harvested and purified by lysing Sf9 cells using successive rounds of sonication and PBS washes. Finally, purified PODS were sterility tested and lyophilized prior to use in experiments. Although equivalence depends on context, 1.5×10^4^ PODS crystals is approximately functionally equivalent to 1 ng of many standard growth factors, cytokines or chemokines in terms of bioactivity (23).

#### Cell culture and differentiation

Monocyte suspension cells (THP-1, Public Health England Culture Collection) were cultured undifferentiated in RPMI-1640 (A10491, Gibco) supplemented with 10% BCS (30-2030, ATCC) (complete medium). For M0 differentiation, THP-1 cells were centrifuged and the conditioned media replaced with fresh complete medium supplemented with 100 ng/ml phorbol 12-myristate-13-acetate (PMA, Sigma P8139). After 48 h, M0 macrophages were further differentiated into M1 or M2 macrophages as described (17,25). Briefly, M0 differentiation medium was removed and adherent M0 cells were washed twice with serum-free medium. For further differentiation, complete medium supplemented with either 100 ng/ml LPS and 20 ng/ml IFN-γ (M1) or 20 ng/ml IL-4 and 20 ng/ml IL-13 (M2) was added to adherent M0 cells and then incubated for up to a further 48h. A375 cells (ATCC) were cultured in DMEM (41966, Gibco) supplemented with 10% FBS (F7524, SIGMA). Cells were seeded at a density of 2e4 cell/cm^2^ and passaged every 3 days. TF-1 cells (ATCC) were cultured in RPMI-1640 supplemented with 2 ng/ml GM-CSF and 10% FBS. Cells were seeded at a density of 2e5 cells/ml and passaged twice a week.

#### Phagocytosis

PODS crystals were either centrifuged onto tissue culture (TC) plates; or centrifuged and dried onto TC plates; or pre-mixed with phagocytic cells for up to 48 hours before plating. Phagocytic cells were then seeded where necessary and subsequently cultured in complete medium. Phagocytosis into cells was demonstrated either by monitoring with a real-time cell history recorder (JuLi stage, NanoEnTek Inc.).

#### Viability assay

To test macrophage viability after PODS uptake, THP-1 cells were seeded at a density of 2e5 cells/ml and differentiated and polarized as described above. 1e6 or 3e6 PODS Empty and PODS FGF-10 per ml were added to unpolarized M0 and polarized M1 and M2 macrophages, resulting in an average of 5 or 15 PODS/ cell, respectively. Cells were then incubated for 48h and 96h hours and a WST-8 assay (Orangu™, Cell Guidance Systems) was performed to determine cell viability.

#### Mobility tracking

THP-1 cells were seeded at a density of 2e5 cells/ml in 24-well plates and differentiated into M0 macrophages as described above. M0 cells were then incubated with PODS IL-2 for 24h in complete medium. The medium was then renewed and cells recorded using a real-time cell history recorder (JuLi stage, NanoEnTek Inc.). Images were taken every 2 min for 24 h and then analyzed using the manual tracker of Image J and the chemotaxis and migration tool (Ibidi).

#### Chemotaxis

THP-1 cells were seeded at a density of 2e5 cells/ml in T25 TC flasks and differentiated into M0 macrophages as described above. M0 cells were then incubated with PODS IL-2 for 24h in complete medium. Subsequently, M0 macrophages were washed with PBS and then detached using an enzyme-free detachment buffer (Cell Dissociation Buffer, Gibco). Cells were then centrifuged, re-counted and seeded in serum-free RPMI-1640 medium at a density of 2e6 cells/ml into the observation area of Chemotaxis µ-Slides (Ibidi). 10% FBS was used as a chemoattractant. Cells in the observation area were recorded using a real-time cell history recorder (JuLi stage, NanoEnTek Inc.). Images were taken once per minute for 12 h and then analyzed using the manual tracker of Image J, and the chemotaxis and migration tool (Ibidi).

#### Directed migration

THP-1 cells were seeded at a density of 2e5 cells/ml in T25 TC flasks and differentiated into M0 macrophages as described above. M0 cells were then incubated with 5e5 PODS eGFP per mL for 24h in complete medium (2.5 PODS/cell). Subsequently, M0 macrophages were washed with PBS and then detached using an enzyme free detachment buffer (Cell Dissociation Buffer, Gibco). Cells were centrifuged, re-counted and seeded in serum-free or 10% FBS containing RPMI-1640 medium at a density of 1e6 cells/ml into 24-well inserts. The bottom well was then filled with either serum-free medium, 10% FBS-containing medium or with 3 day conditioned medium of A375 cells. Microscope images of the bottom well and the insert were taken after 24h of migration.

#### Endogenous IL-6 secretion

THP-1 cells were seeded at a density of 2e5 cells/ml in 24-well plates and differentiated and polarized as described above. The conditioned polarization medium was then collected and stored at -20 °C for later analysis. Macrophages were then incubated with 2e6 PODS FGF-2 per ml for 24h in complete medium (10 PODS/cell). The medium was then collected and stored at -20 °C. Medium samples were tested for the presence of IL-6 by ELISA (DY206, R&D systems) according to the manufacturer’s protocol.

#### Release of IL-6 from PODS IL-6 loaded macrophages

THP-1 cells were seeded at a density of 2e5 cells/ml in 96-well plates and differentiated into M0 macrophages. M0 cells were then incubated with 1e6, 2e6 or 3e6 PODS IL-6 per ml for 24h in complete medium (resulting in 5, 10 or 30 PODS/cell). Cells were then washed twice with PBS and fresh complete medium was added and cells were incubated for 4 days. Additionally, the same number of naked PODS® IL-6 were added to the wells of a 96-well plate and were spun down at 3000 x g for 25 min. The PBS was removed and the plate dried in a laminar flow hood and then supplemented with full growth medium for 4d. After incubation, the medium was collected and subsequently tested by IL-6 ELISA (DY206, R&D systems) according to the manufacturer’s protocol.

#### Functional assay

THP-1 cells were seeded at a density of 2e5 cells/ml in 6-well plates and differentiated and polarized as described above. Macrophages were then incubated with 2e6 PODS FGF-2 per ml for 24h in complete medium (10 PODS/cell). Subsequently, macrophages were washed with PBS and then detached using an enzyme-free detachment buffer (Cell Dissociation Buffer, Gibco). Cells were then centrifuged, re-counted and seeded in serum free TF-1 conditioned medium at a density of 2e6 cells/ml into 24-well inserts. TF-1 cells were seeded at a density of 3e4 cells/cm^2^ in full growth medium into a 24-well plate. After one day of growth, medium was changed to RPMI-1640 supplemented with 0.5% BCS and the macrophage containing inserts were added. Cells were co-incubated for 4 days and the viability of the TF-1 cells was measured performing a WST-8 assay (Orangu™, Cell Guidance Systems).

#### Derivation of primary monocytes from mouse tibia

Murine bone marrow derived monocytes were isolated according to Wagner et al(16). Briefly, the tibias of 3 C57BL/6 mice were prepared for harvest. After washing once in 96% ethanol and twice in PBS, the distal end of each bone was cut with fine scissors and the bones flushed with warm medium (M199 supplemented with 10% FBS and 1% Penicillin/Streptomycin) using a 28-G needle and a 1 ml syringe. The flow-through from the bones was collected and filtered through a 70 µm cell strainer. The cell suspension was centrifuged at 200 x g for 10 min at RT. The pellet was washed with 25 ml of medium and centrifuged again. The cells were then seeded at 0.5e6 cells/ml in 6-well ultra-low-attachment plates. Cells were cultured for 5 days before using in experiments.

## Discussion

Here, we have explored the potential utility of PODS protein crystal nanoparticles as vectors for modifying the cytokine secretion profile of phagocytic cells. We first demonstrated that PODS are readily, consistently and efficiently taken up by professional phagocytes including murine bone marrow-derived monocytes and THP-1 cell line-derived macrophages regardless of polarization states.

A potential problem with protein nanoparticle loading into macrophages is destruction of the particle and any cargo by the phagolysosome due to its low pH and lysosomal proteases. PODS are particularly stable in acidic environments and protect their cargo due to their tightly packaged format (26). Following phagocytosis, we have demonstrated sustained release from macrophages of a cytokine encapsulated into PODS crystals. The released cytokine could be detected reliably, and in a dose-dependent manner, in macrophage culture medium. This was observed with macrophages averaging as few as 5 PODS per macrophage. Even more importantly, released protein was shown to be bioactive as verified by proliferation of an FGF-2 reactive cell line upon co-cultured with PODS FGF-2 loaded macrophages.

It isn’t clear how PODS crystals are able to withstand the internal machinery of macrophages and secrete their cargo intact. One possibility is that the unique structure of PODS allows them to endure, or simply overwhelm, the hostile phagolysosome environment. However, work by others suggests an alternative mechanism. Detection of active cargo released from macrophages was also demonstrated for macrophages loaded with antiretroviral nanoparticles (27) which avoided intracellular degradation and were recycled to the plasma membrane. The perinuclear localization of PODS crystals after uptake that we observed for PODS crystals would support such a mechanism.

Importantly, the uptake of PODS doesn’t seem to be cytotoxic nor does it fundamentally change characteristics of professional phagocytes such as mobility or chemotaxis. This lack of cytotoxicity is seen even when multiple PODS crystals are loaded into a single macrophage swelling the cell’s size. Maintenance of functionality is an important pre-requisite for most useful applications of any macrophage nanoparticles.

Selective targeting of drugs to diseased tissue, notably cancer, is urgently needed to reduce levels of off-target toxicity and to increase efficacy. The use of cells as a therapeutic delivery system has been under discussion since the 1970s (28) and the use of macrophages for a Trojan horse drug delivery strategy to treat cancer was proposed in the 1990s (29).

Macrophages are particularly attractive for cancer therapy because of their tumor infiltrating behavior. Most macrophages in cancer are derived from haemopoietic lineages and are continuously recycled with a lifespan within the cancer of about two weeks. Developing an effective particle which contains the drug, allows function of the macrophage and subsequent drug release has been challenging, particularly for protein drugs such as cytokines. Despite ongoing research there has been no cell-based delivery system approved for the clinic, although two therapeutics using red blood cells as drug vehicle are currently in phase III clinical trials (30,31).

Efficient uptake of protein nanoparticles by phagocytes may allow the use of PODS crystals in a Trojan horse strategy to deliver these cytokine depots preferentially or specifically to an area of disease by exploiting the tumor infiltrating behavior of macrophages (32). In order for a PODS Trojan horse approach to be viable as a delivery tool for therapeutic proteins, several criteria have to be met. A basic requirement for a drug delivery system is that it does not produce systemic inflammation or cytotoxicity. PODS proteins have already been used to deliver a diverse range of cargo proteins in different animal models and no adverse effects were observed, including in studies of bone remodeling in rats (23) and dogs (33); as well as neuronal and cartilage regeneration in mice (manuscripts in preparation). While there were no signs of increased inflammation detectable in these animal models which included examinations of distant sites such as the lung, a detailed investigation into interactions between macrophages and PODS and their possible cytotoxic effects remains to be conducted.

Our studies showed that PODS uptake did not impair three attributes required for macrophage tumor infiltration: mobility, chemotaxis and migration through narrow spaces. Our results demonstrated no effect on overall levels of mobility. The PODS-loaded macrophages could also squeeze through narrow capillary like tubes and migrate towards a chemotactic signal whether that was towards sera or towards chemotactic agents secreted by cancer cells. Therefore, our results indicate that the tumor homing ability of macrophages may not be significantly affected by phagocytosis of PODS or by the embedded IL-2 cargo protein.

The cytokine cargos of PODS-loaded macrophages may be used to shift the tumor microenvironment in several ways. Macrophages can switch between M1 or an M2 phenotypes by responding to cues from the local environment. However, this polarization can be transient and is non-binary with high levels of cell plasticity (34). *In vitro*, the secretion of the cytokine IL-6 is one of many markers used to determine the polarization status of M1 macrophages (35,36) and was shown to be dependent on particle size, with the highest secretion seen with particles of 0.8 μm (37).

FGF-2 has been shown to shift macrophages towards the M2-like phenotype (38). A shift in polarization could be exploited in cancer therapy. Tumor-associated macrophages (TAMs) usually express an M2 phenotype (alternatively activated subset), which leads to them performing immunosuppressive and tumor promoting functions. Reprogramming of such cells towards an M1-like phenotype (classically activated subset) may suppress their pro-cancer phenotype and unleash anti-tumor activity (39,40).

Cytokines which reprogram the activity of other immune cells could also be used as cargo. Cytokines are key regulators of immune cell activity and there are many potential targets and mechanisms with the TME which could be addressed. One obvious candidate is IL-2 which is approved for the treatment of metastatic renal cell cancer and metastatic melanoma (41).

The use of high doses is required to achieve efficacy which result in high levels of toxicity requiring hospitalization during therapy. In particular, IL-2 causes leaky vasculature (42). Targeting IL-2 to tumors using macrophages would shift the therapeutic window to increase efficacy whilst reducing toxicity and allow IL-2 therapy to become more widely used.

This study lays the foundations for the development of PODS for use in phagocytic cell research in general and macrophages in particular. It also provides a proof-of-concept for a PODS-based macrophage mediated Trojan horse strategy for delivering proteins including cytokines to combat cancer and other diseases.

## Acknowledgements

The authors would like to thank Dr Ciara Whitty and Dr Raj Gandhi for early observations and initial characterizations of interactions of PODS with non-professional phagocytes. Furthermore, we would like to thank Prof Ann Ager for helpful discussions and comments on the manuscript.

## Author contributions

AW and NJ collected and assembled the data.

AW, CP, MJ designed the study, analysed and interpretated the data and prepared the manuscript.

All authors provided final approval of the article prior to submission.

## Figure captions

**Supp fig 1: PODS morphology**. PODS Empty were spun down onto tissue culture plastic and imaged using SEM (left image) and brightfield microscopy (right image) at a magnification of 40x. PODS are easily distinguishable from cells due to their cubic shape.

**Supp video 1: Mobility of BMDMs after PODS GM-CSF uptake**. BMDMs loaded with an average of 5 PODS GM-CSF per cell were monitored with a life cell imaging system for 24h. This video shows 52s equivalent to approximately 4 h and 20 min in real time.

**Supp fig 2:** GM-CSF loaded mouse BMDM were incubated with PODS eGFP for 24 h and monitored using a live cell imaging system at 1 frame/min. The dashed white circle follows the movement of a macrophage towards a single PODS, the uptake of the PODS, and the contraction of the macrophage over 21 min depicted in a still of 4 frames (59, 70, 76 and 80).

**Supp video 2: Uptake of PODS eGFP into PODS GM-CSF loaded BMDMs**. BMDMs loaded with an average of 5 PODS GM-CSF per cell were incubated with PODS eGFP at a ration of 5 PODS eGFP per cell and monitored with a life cell imaging system for 24h. This video shows 26s equivalent to approximately 2 h and 10 min in real time.

**Supp Fig 3: Efficient uptake of PODS into M0**. THP-1 cells were differentiated first into M0 macrophages and then further polarized into M1 cells. Uptake of PODS Empty was allowed for 24h and monitored using a live cell imaging system at 1 frame/min. Four frames (1, 675 and 1335) are shown to illustrate the uptake over time.

**Supp Fig 4: Phagocytosis of PODS protein crystals into non-professional phagocytes**. Chondrocytes, NIH3T3 cells and C2C12 cells were cultured with PODS Empty for 24 h and subsequently imaged with brightfield microscopy at either 20x or 40x magnification. The dashed square marks the area of the zoom shown in the right image of each panel.

**Supp Fig 5: Viability of PODS-loaded macrophages after 96h**. M0, M1 and M2 macrophages were incubated with PODS Empty or PODS FGF-10 (1:5 and 1:15) for 24 h in a 96-well TC plate. Subsequently, medium was changed and viability of cells was measured 96h after PODS uptake using a colorimetric assay (Orangu™, Cell Guidance Systems). The fold change in viability was calculated relative to unloaded macrophages.

**Supp Fig 6: Overloading of macrophages leads to apoptosis**. M1 cells already loaded with an average of 10 PODS per cell, were incubated with another 10 PODS per cell for further 24h and images were taken every 2 min. Three frames were chosen to follow 3 cells. The dashed white circle marks a healthy cell with approximately 10 PODS ingested. The white arrow and turquois arrow mark two cells with more than 50 PODS ingested; both cells undergo apoptosis shown in frame 190 and frame 690 respectively.

