## Supplementary figures and images for "Phagocytosed polyhedrin-cytokine co-crystal nanoparticles provide sustained secretion of bioactive cytokines from macrophages"

### Supplemental Figure 1

#### Figure S1

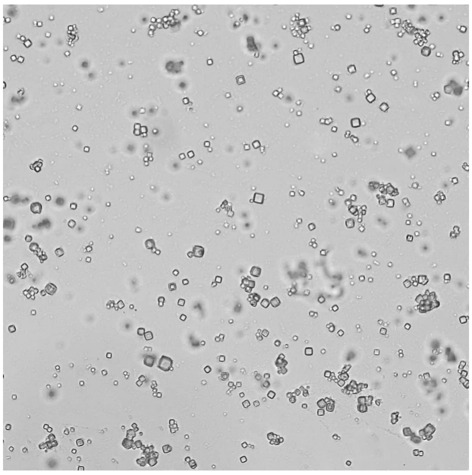

Brightfield 40x

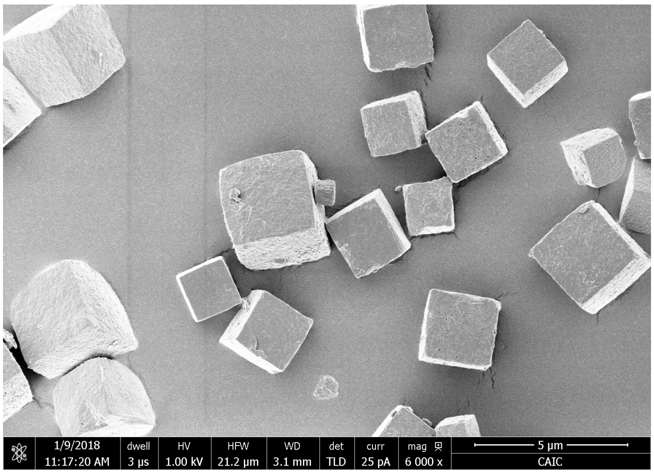


SEM

### Supplemental Figure 2

#### Figure S2

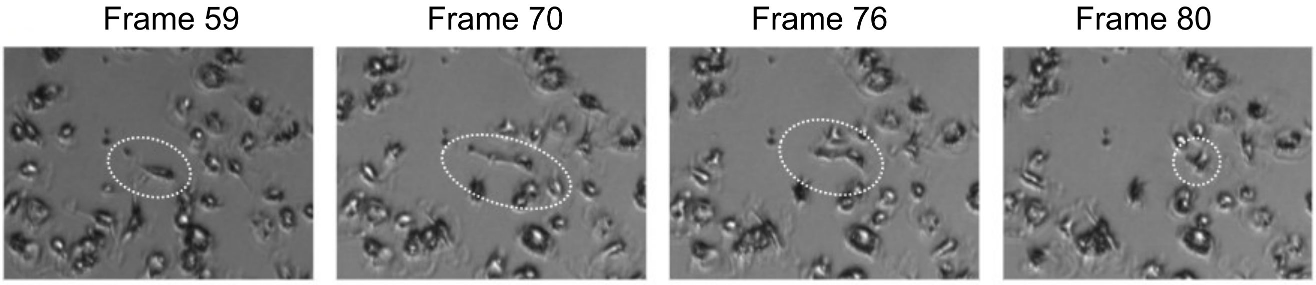

### Supplemental Figure 3

#### Figure S3

Frame 1 | Frame 675 | Frame 1335

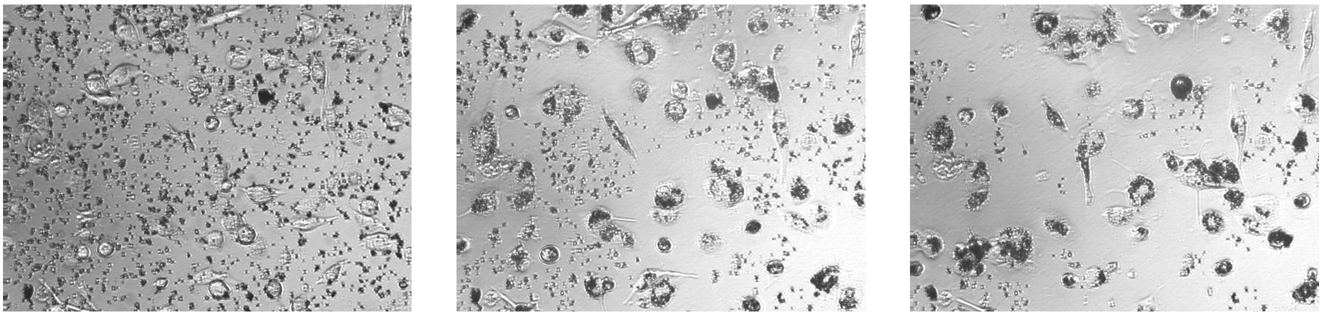

### Supplemental Figure 4

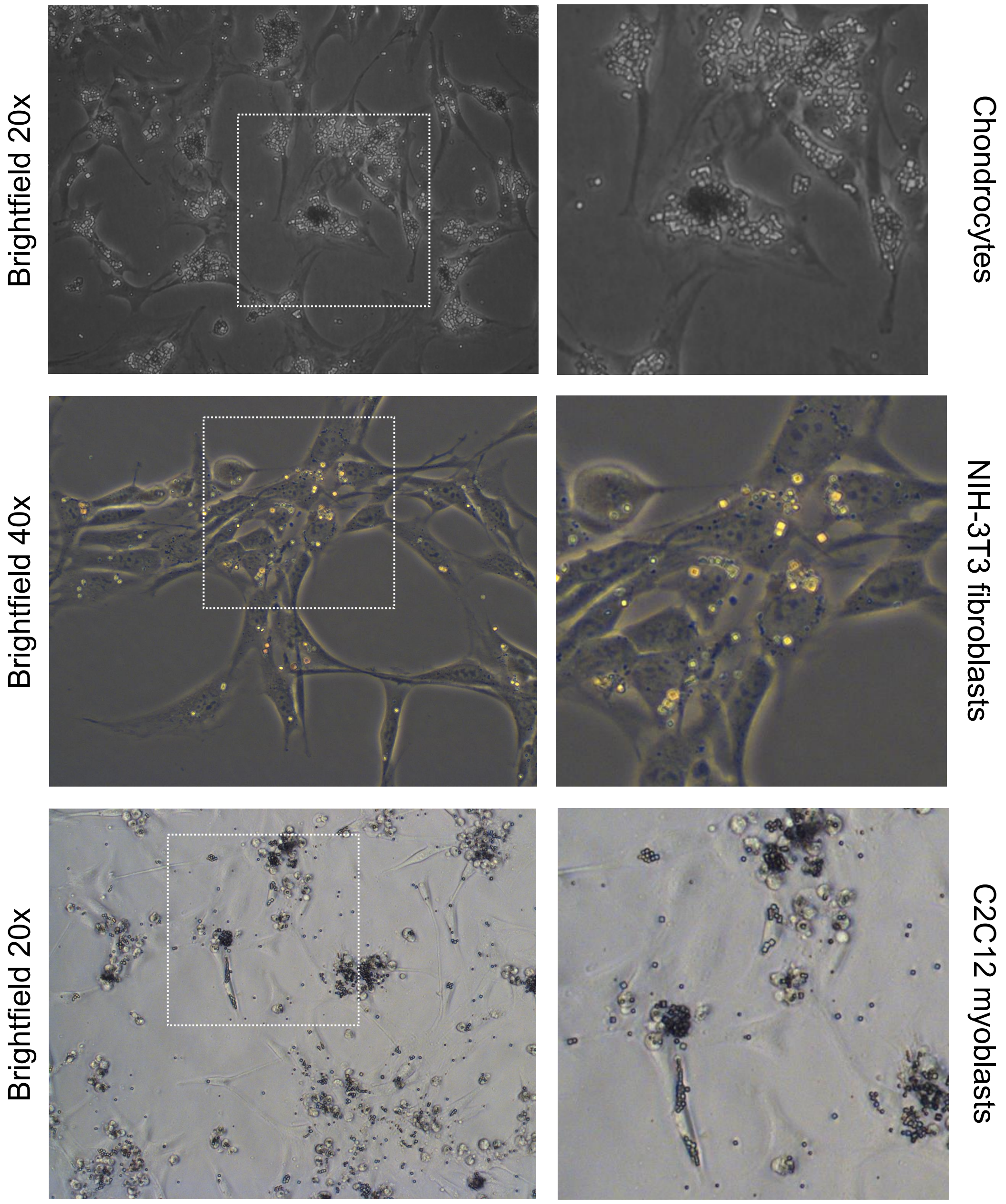

### Supplemental Figure 5

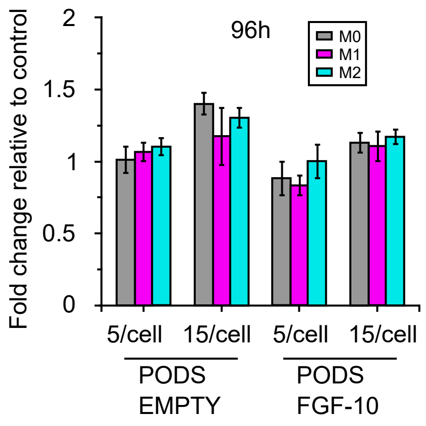

### Supplemental Figure 6

#### Figure S6

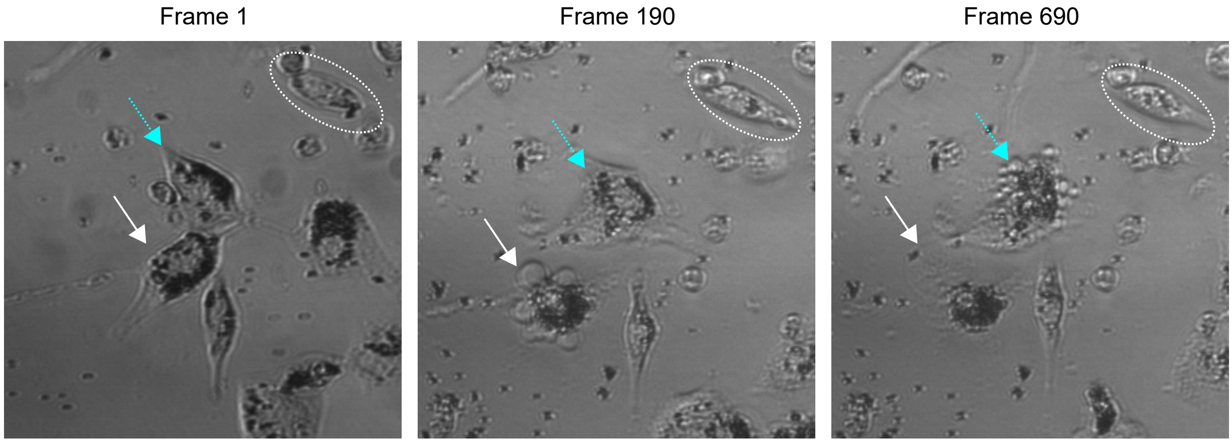
